# Voxel-wise or Region-wise Nuisance Regression for Functional Connectivity Analyses: Does it matter?

**DOI:** 10.1101/2024.12.10.627766

**Authors:** Tobias Muganga, Leonard Sasse, Daouia I. Larabi, Nicolás Nieto, Julian Caspers, Simon B. Eickhoff, Kaustubh R. Patil

## Abstract

Removal of nuisance signals (such as motion) from the BOLD time series is an important aspect of preprocessing to obtain meaningful resting-state functional connectivity (rs-FC). The nuisance signals are commonly removed using denoising procedures at the finest resolution, i.e. the voxel time series. Typically the voxel-wise time series are then aggregated into predefined regions or parcels to obtain a rs-FC matrix as the correlation between pairs of regional time series. Computational efficiency can be improved by denoising the aggregated regional time series instead of the voxel time series. However, a comprehensive comparison of the effects of denoising on these two resolutions is missing.

In this study, we systematically investigate the effects of denoising at different time series resolutions (voxel- and region-level) in 370 unrelated subjects from the HCP-YA dataset. Alongside the time series resolution, we considered additional factors such as aggregation method (*Mean* and *first eigenvariate* [*EV*]) and parcellation granularity (100, 400, and 1,000 regions). To assess the effect of those choices on the utility of the resulting whole-brain rs-FC, we evaluated the individual specificity (fingerprinting) and the capacity to predict age and three cognitive scores.

Our findings show generally equal or better performance for region-level denoising with notable differences depending on the aggregation method. Using mean aggregation yielded equal individual specificity and prediction performance for voxel- and region-level denoising. When EV was employed for aggregation, the individual specificity of voxel-level denoising was reduced compared to region-level denoising. Increasing parcellation granularity generally improved individual specificity. For the prediction of age and cognitive test scores, only fluid intelligence indicated worse performance for voxel-level denoising in the case of aggregating with the *EV*.

Based on these results, we recommend the adoption of region-level denoising for brain-behavior investigations when using mean aggregation. This approach offers equal individual specificity and prediction capacity with reduced computational resources for the analysis of rs-FC patterns.

## Introduction

In recent years, resting-state functional connectivity (rs-FC) has received increased interest as a representation of individual-level functional brain organization that is predictive of behavior and potentially suitable as a biomarker (Dubois & Adolphs, 2016; Finn & Rosenberg, 2021). The rs-FC refers to the statistical relationships between spontaneous fluctuations in the blood oxygen level-dependent (BOLD) signal of different brain regions, typically measured with functional magnetic resonance imaging (fMRI), while a subject is at rest (B. Biswal et al., 1995; Friston, 2011). For rs-FC to be suitable as an individual-level biomarker, it must be predictive of inter-individual differences and be stable for repeated measurements of the same individual (Finn & Rosenberg, 2021; Gordon et al., 2017; Mantwill et al., 2022). Studies have found patterns in rs-FC to be individual-specific (Amico & Goñi, 2018; Finn et al., 2015) and stable over extended periods (Gratton et al., 2018; Horien et al., 2019). Additionally, rs-FC patterns have been linked to individual differences in cognitive, behavioral, and clinical attributes (Finn & Constable, 2016; Geerligs et al., 2015; Kong et al., 2019; Yamada et al., 2017).

However, capturing meaningful whole-brain rs-FC patterns at the individual level is challenging, as the measured BOLD signal in each voxel is a mixture of neuronal and non-neuronal nuisance signals (Bianciardi et al., 2009; B. B. Biswal et al., 2010; Fox & Raichle, 2007; Logothetis, 2003; Power et al., 2015). Reducing the influence of such nuisance signals is commonly done by removing their variance from the BOLD time series using linear regression (Power et al., 2014). Such removal methods are commonly applied to each voxel’s time series, i.e. the finest resolution, which can be computationally expensive. As rs-FC is calculated using aggregated regional time series, applying the removal process to the regional time series, instead of the voxel time series, offers a potential reduction in computational resources. However, the effect of this choice on the rs-FC and its utility in individual-level analysis has not been explored. Therefore, moving nuisance removal to the regional level requires additional considerations regarding the data quality and the region definition and how it affects the individual-specificity and predictability of behavior (Airan et al., 2016; Eickhoff et al., 2018; Finn & Rosenberg, 2021; Power et al., 2015; Siegel et al., 2017).

Methods applied on the voxel-wise time series to increase the data quality (i.e. denoising) frequently include removing slow signal drifts, filtering the frequency bands to spectra associated with neuronal signals (< 0.1 Hz), and regressing nuisance estimates (B. Biswal et al., 1995; Leopold et al., 2003; Power et al., 2015). Several nuisance estimates can be derived from different origins; parameters from spatial realignment due to subjects head motion are often expanded (Friston et al., 1996; Satterthwaite et al., 2013), or signals from non-grey matter are used (Behzadi et al., 2007). Unfortunately, head motion affects voxels nonuniformly across the brain (Power et al., 2017). Therefore, even after using more powerful data-driven estimation of noise components like independent component analyses (ICA; Ciric et al., 2017; Parkes et al., 2018; Salimi-Khorshidi et al., 2014) some motion-related nuisance variance might remain (Bright et al., 2017; Kopal et al., 2020; Power et al., 2015). Performing the removal process at the aggregated regional time series might be advantageous for stabilizing not only the neuronal signal (Korhonen et al., 2017) but also the motion-related nuisance signal. This stabilized nuisance signal might better match the nuisance estimates, enabling more efficient and effective removal. However, this potential rests on choices of functionally coherent regional representation to enable stable and meaningful interpretability (Airan et al., 2016; Eickhoff et al., 2018; Power et al., 2015).

In order to generate a functionally coherent regional time series the voxels within a region should exhibit homogeneous functional activity (Stanley et al., 2013). Conversely, functional heterogeneity can arise when multiple non-related functional units are captured by the same region definition (Friston et al., 2006; Korhonen et al., 2017). Heterogeneity can arise due to a subject’s misalignment to the parcellation or coarse granularities, i.e. large regions (Craddock et al., 2012). This makes the selection of an aggregation method critical as its behavior in homogeneous and heterogeneous regions will impact the quality of the rs-FC and in turn the individual differences captured by rs-FC. A common approach to aggregate voxel-wise time series into homogeneous regional time series is averaging all voxels’ time series in a region, thereby capturing shared aspects by effectively smoothing the time series and reducing complexity (Korhonen et al., 2017). While straightforward to apply and interpret, it bears the risk of suppressing or canceling relevant variance if the region spans multiple functionally distinct units (Friston et al., 2006). An alternative approach is computing the first eigenvariate (EV; (Friston et al., 2006). It results from performing eigendecomposition on the voxel time series, resulting in a weighted mean reflecting the main variance component of the regional response (Saxe et al., 2006). In a functionally homogeneous region, the mean and EV time series are similar (Airan et al., 2016; Saxe et al., 2006). Conversely, in a functionally heterogeneous region, the EV will select only the most dominant part of the regional variance (Saxe et al., 2006). The EV’s sensitivity can lead to regional time series dominated by a subset of the voxels within the region that might differ from subject to subject (Craddock et al., 2012; Saxe et al., 2006). In the context of individual differences, the EV’s sensitivity might be desirable to account for small subject-specific misalignments to region definitions which are often derived from a group-level atlas. Whether this characteristic is advantageous in the context of the analysis of individual differences is still unclear.

In this regard, the granularity of the parcellation scheme has a considerable impact as the differing sizes of regions affect the functional homogeneity of the regions (Craddock et al., 2012; Shen et al., 2013). With larger regions more smoothing is applied which might lead to loss of individual differences (Airan et al., 2016; Finn et al., 2015; Korhonen et al., 2017). On the other hand, a highly granular parcellation might be too close to voxel-level resolution, overemphasizing the impact of residual motion per region, promoting spurious relationships (Power et al., 2015; Stanley et al., 2013) and increasing computational complexity. However, there is no single correct number of regions, as different granularities reflect distinct organizational layers of the brain (Eickhoff et al., 2018), which might impact the stability and predictiveness of individual differences nonuniformly.

To our knowledge, no previous work has investigated the effects of region-wise denoising for individual differences analyses. While for mean aggregated regional time series, both voxel-wise and region-wise denoising should result in the same rs-FC patterns at any granularity (Friston et al., 2006), the behavior of EV aggregation remains unexplored. To fill this gap, we systematically evaluated the effects of denoising the voxel-level and region-level time series. We assessed the individual specificity and predictability of individual differences in resulting rs-FC (Finn & Rosenberg, 2021). Specifically, we evaluated individual specificity as the ability to identify the same subject from different scans with the binary approach of connectome fingerprinting (Finn et al., 2015) and the continuous approach of differential identifiability (Amico & Goñi, 2018). The behavioral predictability was addressed by predictive analysis of age and cognitive test scores acquired outside the scanner. We included considerations to test how known influences such as the quality of the data and parcellation granularity influence region-level denoising. To this end, we evaluated the same dataset at two levels of preprocessing affecting data quality and assessed the association of obtained rs-FC with motion, and computed rs-FC with different granularities of the same parcellation scheme. Our comprehensive analyses can inform the design of fMRI processing pipelines and help save computational resources.

## Methods

### Dataset

The data used for this study was obtained from the Human Connectome Project’s Young Adult (HCP-YA) S1200 subject release (Van Essen et al., 2013). The HCP-YA dataset includes two resting-state sessions (R1 and R2) carried out on consecutive days with 30 minutes of resting-state acquisition per day. Each session involved 2 × 15-minute runs with opposite phase encoding directions (left-right [LR] and right-left [RL]). All subjects’ resting-state scans were acquired with a 3T Siemens scanner using the same protocol: slice-accelerated multiband pulse sequence (factor of 8), a spatial resolution of 2 mm isotropic voxels, TR of 720 ms, TE of 33 ms, with a total of 72 slices, a field of view of 208 mm in the anterior-posterior direction, 180 mm in the phase encoding direction (LR and RL), and 144 mm in the inferior-superior direction (for in-depth details on the acquisition protocol, see Glasser et al., 2013; Uğurbil et al., 2013; Van Essen et al., 2013). The HCP-YA scanning protocol was approved by the local Institutional Review Board at Washington University in St. Louis, and all participants provided written consent before participation. Approval for retrospective analysis was given by the ethics committee at the Faculty of Medicine at Heinrich-Heine University Düsseldorf (Study No. 2018-317-RetroDeuA). For our investigation, 370 (178 female) non-related individuals were selected (by Family_ID) for whom cognitive scores and rs-fMRI scans from both resting-state sessions (R1 and R2), acquired with both phase-encoding directions (LR and RL) were available (Table 1).

**Table 1.** Basic summary statistics for demographics and cognitive variables. Original variable names specified by the HCP are provided in parentheses.

| Variable | Statistic |
| --- | --- |
| Sex, % (N) | Male: 52% (192)<br>Female: 48% (178) |
| Age, M (SD) | 28.8 (3.8) |
| Working memory (ListSort_Unadj), M (SD) | 111.8 (11.8) |
| Fluid intelligence (PMAT24_A_CR), M(SD) | 16.9 (4.8) |
| Reading (ReadEng_Unadj), M (SD) | 117.1 (10.8) |
| Motion (mean FD), M (SD) | 0.16 (0.05) |

#### HCP preprocessing

The HCP-YA resting-state data is accessible (www.dbconnectome.com) at different points throughout the HCP processing pipelines. The preprocessing of the HCP-YA dataset has been described extensively elsewhere (Glasser et al., 2013; Smith et al., 2013). In short, the minimal preprocessing pipeline applied by the HCP included the correction of spatial distortion, realignment to correct for head motion to a single-band reference image, distortion field correction, and registration to 2 mm Montreal Neurological Institute (MNI) standard space (MNI152NLIN6Asym). The necessary estimates per step were separately calculated, concatenated, and applied to the fMRI volumes in one spline interpolation to minimize interpolation-induced blurring. The resulting time series were additionally intensity normalized (to a global mean of 10,000) and brain masked (described in detail by Glasser et al., 2013).

The ICA-FIX (FMRIB’s ICA-based X-noiseifier; Salimi-Khorshidi et al., 2014) denoising pipeline used the minimally preprocessed data as input and removed non-neuronal components from each voxel’s time series. This pipeline included temporal high-pass filtering (cutoff at 2,000 s full-width-half-minimum), FIX classification of 250 independent components (from ICA) into “good” and “bad” components, and regression of “bad” components and 24 regressors derived from motion estimation (Smith et al., 2013).

For our analyses, we used the minimally preprocessed data and the data from the ICA-FIX-based denoising pipeline (Smith et al., 2013). This enabled us to do an assessment of the influence of the amount of denoising performed. The two data quality configurations will be respectively referenced as *Minimal* and *ICA-FIX* from here on forward.

### Rs-FC processing

Our rs-FC processing pipelines (Figure 1) considered two main aspects: denoising and parcellation. After downloading a given subject’s volume, voxel-wise time series were extracted with a binarized brain mask generated from the parcellation template using the nilearn toolbox (Abraham et al., 2014). Next, either denoising or parcellation was performed. As a last step in each pipeline, rs-FC was generated by calculating the Pearson’s correlation coefficient between all pairs of z-scored regional time series (Pernet et al., 2013) resulting in a N_Region_ × N_Region_ correlation matrix. Each subject’s rs-FC was computed for each phase-encoding run separately.

**Figure 1.**
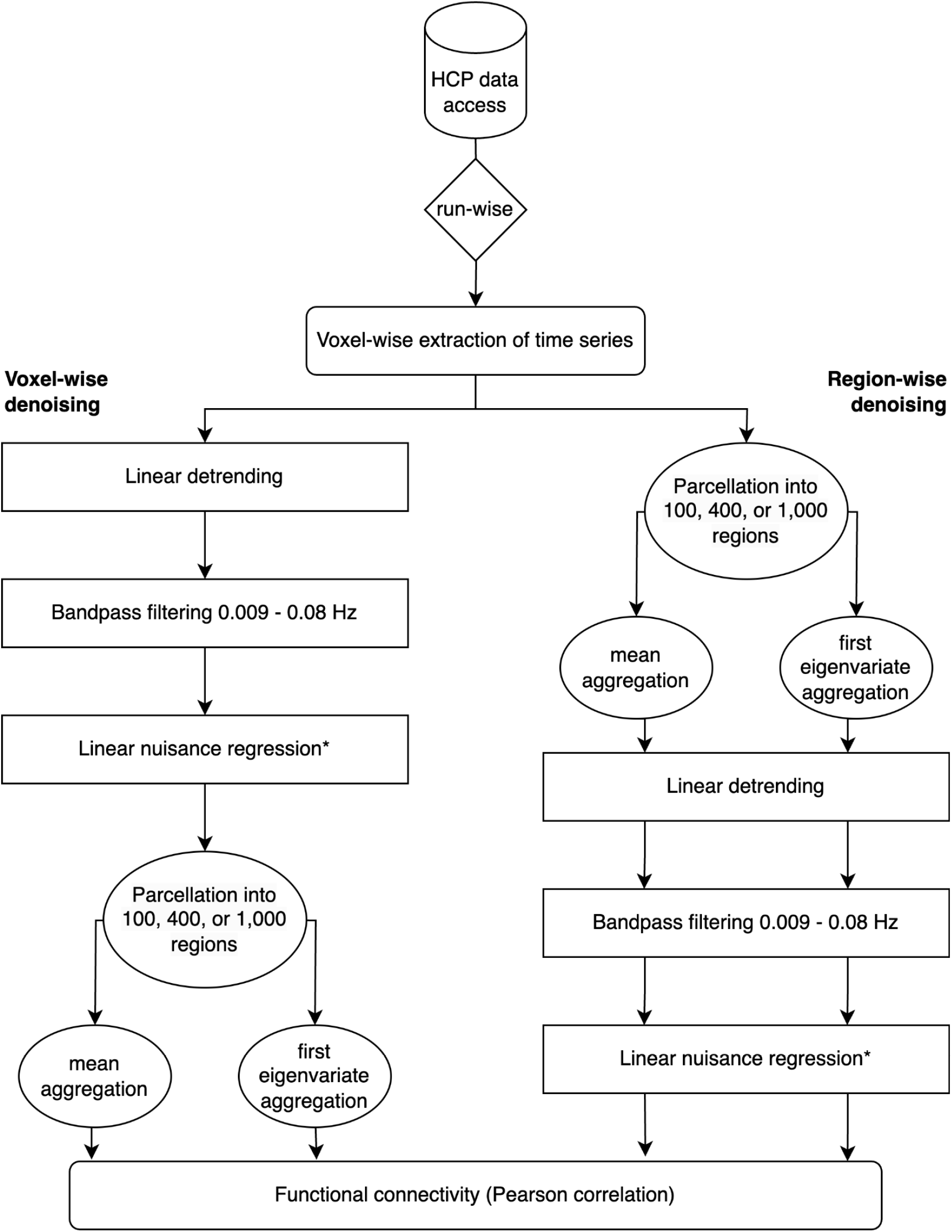
The schematics illustrate the processing pipelines for voxel-wise and region-wise denoising. Following the download, each phase encoding run is processed independently. Voxel-wise time series extraction precedes either denoising or parcellation. The regressors include WM, CSF, GS, their temporal derivatives, and square terms. *24 motion regressors were included for the minimally preprocessed data configuration only.

#### Voxel-wise and region-wise denoising

To evaluate differences between denoising the BOLD time series voxel-wise and region-wise we set up pipelines that differed in two main aspects. Switching the sequence in which these two sections were applied led to the generation of either rs-FC denoised at the voxel level or at the region level. The voxel-wise pipeline (Fig. 1, left) denoised all the extracted voxel time series, and aggregated the denoised voxel time series into regional time series. The region-wise pipeline (Fig. 1, right) aggregated the voxel-wise extracted time series into regional time series first, reducing dimensionality, and applied denoising to the regional time series. In either case, rs-FC was calculated using regional time series that had either been denoised voxel-wise or region-wise.

### Denoising parameters

Denoising included the removal of linear trends to correct gradual signal drifts over time, Butterworth bandpass filtering in the frequencies between 0.009 and 0.08 Hz (Kopal et al., 2020; J. Li et al., 2019; Siegel et al., 2017), and linear nuisance regression. All of these steps were performed using nilearn’s “signal.clean” function. A total of 12 nuisance regressors provided by HCP were used consisting of mean signals generated from white matter (WM) and cerebral spinal fluid (CSF) masks as well as the mean global signal (Fox et al., 2009), their temporal derivatives, and their squared terms (Satterthwaite et al., 2013). While effective at reducing noise, removing GS is heavily debated as it may also remove signals of interest, potentially distorting individual differences (Demertzi et al., 2022; Fox et al., 2009; J. Li et al., 2019; Macey et al., 2004). However, we decided to include it since removing GS can improve the predictability of behavior from rs-FC (J. Li et al., 2019; Yan et al., 2013). Additional 24 motion regressors (3 rotations, 3 translations, their 6 temporal derivatives, and 12 squared terms) were included for denoising of the *Minimal* data configuration as HCP’s ICA-FIX-based denoising pipeline already removed 24 additional motion regressors (Smith et al., 2013). To prevent the reintroduction of artifacts due to the modularity of our setup nilearn orthogonalizes regressors to the bandpass filter before removal (Lindquist et al., 2019).

#### Generation of regional time series

### Aggregation Metrics

Two aggregation metrics were employed to evaluate the ability to capture regional time series capturing individual differences at different levels of processing. The aggregation into regional time series was either performed on the denoised or the voxel time series. To generate regional time series, all voxel-wise time series from a region were aggregated into one regional time series by computing the *Mean* (Stanley et al., 2013) or the *EV* (Friston et al., 2006). To generate the *EV* the voxel-wise time series were decomposed using single-value decomposition and the resulting first eigenvector is sign adjusted and scaled by the square root of the variance it explains.

### Whole brain parcellation

Previous studies suggest that parcellation granularity (i.e., region size) contributes to the ability to identify individual differences (Airan et al., 2016; Finn et al., 2015). Therefore, we generated rs-FC using three different granularities (100, 400, and 1,000 regions) of the Schaefer parcellation scheme. The granularities represent the coarsest and finest granularities provided as well as a commonly used intermediary described by Schaefer et al. (2018) to capture fully differentiated regions.

The Schaefer parcellation templates were used (https://github.com/ThomasYeoLab/CBIG/tree/master/stable_projects/brain_parcellati on/Schaefer2018_LocalGlobal) in the Yeo 7-network version in MNI152 (MNI152NLIN6Asym) space (Grabner et al., 2006) to match the data provided by the HCP-YA. This parcellation scheme is based on the resting-state and task-fMRI data of 1,489 subjects. Scans between subjects were aligned with surface-based registration, and cortical regions ranging from 100 to 1000 (in steps of 100) were identified with a gradient-weighted Markov Random Field model. The parcellation generated by this approach has been found to produce parcels containing functionally homogeneous voxel time series and with neurobiologically meaningful boundaries, i.e., they agree well with histologically defined boundaries. The parcellation template was used to mask the voxel-wise extracted time series, assigning labels to each voxel time series. Subsequently, voxel time series with the same label according to the parcellation scheme were aggregated into a regional time series. Subcortical regions were not included in the analysis.

### Stability of individual specificity of rs-FC: Identification Accuracy & Differential Identifiability

We assessed the stability of rs-FC in two ways: identification accuracy (**I_acc_**; (Finn et al., 2015) and differential identifiability (**I_diff_**; (Amico & Goñi, 2018). First, the rs-FC matrices from the different phase-encoding runs were averaged to arrive at one scan per session resulting in two rs-FC per subject (R1 and R2). Next, we created a N_subject_ × N_subject_ identifiability matrix by calculating Spearman correlations between the rs-FC of the two sessions (R1 and R2). In this identifiability matrix, the within-subject correlations are represented by the diagonal values, and the between-subject correlations are represented by off-diagonal values. Identification accuracy was then calculated as the proportion of subjects for whom the highest correlation value was positioned on the diagonal. Differential identifiability was calculated by subtracting the average between-subject correlations from the average within-subject correlations.

### Prediction of age and cognitive scores

To test the effects of denoising the time series voxel- or region-wise on brain-behavior predictions, we used the rs-FC obtained from the different preprocessing pipelines to predict age as well as three cognitive measures previously demonstrated to result in reasonable prediction accuracy in the HCP-YA dataset (He et al., 2020; Sasse et al., 2023): working memory (ListSort_Unadj), fluid intelligence (PMAT24_A_CR), and reading (ReadEng_Unadj). First, each subject’s four rs-FC matrices were averaged across phase encoding directions (LR and RL) and sessions (R1 and R2) to generate one rs-FC per subject. The unique edges from this rs-FC matrix (i.e., the lower triangle) were then used as features. A kernel ridge regression model with Pearson kernel was used within a nested 5-fold cross-validation (CV) scheme with 5 repetitions where the inner 5-fold CV was used to select the l2-regularization parameter. To control for possible confounding influences, we removed variance associated with age, sex, and motion (i.e., framewise displacement [FD]; He et al., 2020) from the cognitive scores in a CV-consistent manner, i.e. confound regression models were trained on the training data and applied to both training and test data to prevent data leakage (More et al., 2023; Snoek et al., 2019). The prediction accuracy was assessed as Pearson’s correlation between the predicted value and the observed value of the target. The prediction pipeline was implemented in Python (Version 3.10.0) using the Julearn package (Version 0.2.6.dev3; Hamdan et al., 2023). Julearn is an extension built on top of scikit-learn (Version 1.1.3; (Pedregosa et al., 2011).

### Motion association assessment for rs-FC

To understand how the main results might be impacted by residual head motion influences, we also computed rs-FC typicality (TFC; Kopal et al., 2020). TFC evaluates the typicality of an individual rs-FC compared with a group-averaged rs-FC. A higher typicality is associated with lower motion estimates (i.e. mean FD). We used this measure to assess the motion association of rs-FC resulting from our different pipelines. TFC was computed following rs-FC generation for each run separately. TFC values were generated by computing Pearson’s correlation of rs-FC to the group rs-FC of the same run. The group rs-FC was generated by averaging all rs-FC from the same run. The association between TFC and FD was assessed by computing the Spearman correlation.

### Preprocessing time assessment

The computation time of each pipeline execution was tracked to compare the runtime. Once the data download was complete, the timer was started before voxel-wise time series extraction and stopped after the generation of rs-FC. To ensure the stability of the measurements per subject, we made sure that all pipelines for a single subject were executed on the same compute node on a high-throughput computing cluster. Although different subjects might have been processed with slightly variable processing power, the speed is stable across all pipeline executions for each subject.

### Data and Code availability

Further details regarding the obtaining of the HCP-YA dataset can be found at: https://www.humanconnectome.org/

The preprocessing, analysis, and visualization scripts used in this project are available in the project repository: https://github.com/juaml/voxel-vs-region-denoising

The preprocessing pipeline utilized the open-source toolbox **Junifer**, which is accessible at: https://github.com/juaml/junifer

Machine learning-based prediction analyses were conducted using the toolbox

**Julearn**, available at: https://github.com/juaml/julearn

## Results

### Individual-specificity of rs-FC denoised at the voxel- or region-level

Consistent with previous literature using whole-brain rs-FC (Finn et al., 2015; Sasse et al., 2023), ***I_acc_*** for voxel-wise denoising ranged from 0.768 to 0.997, while region-wise denoising ranged from 0.945 to 0.997 (Table 2). Region-wise denoising was equal or better in identification performance than voxel-wise denoising in all the cases. For ***I_diff_***, the performance for both voxel-wise denoising and region-wise denoising ranged from 19.58 to 34.768 (Table 2). For this metric, the region-wise denoising also obtained better or equal results in all cases except the *EV* aggregation with a parcellation granularity of 100 regions.

**Table 2.** Overview of Identification performance across all conditions.

| Row | Data configuration | Parcellation granularity | $I_{acc}$<br>(Voxel) | $I_{acc}$<br>(Region) | $I_{diff}$<br>(Voxel) | $I_{diff}$<br>(Region) |
| --- | --- | --- | --- | --- | --- | --- |
| Aggregation: *EV* |  |  |  |  |  |  |
| 1 | Minimal | 100 | 0.802 | 0.951 | 22.37 | 19.67 |
| 2 | Minimal | 400 | 0.959 | 0.986 | 27.954 | 28.041 |
| 3 | Minimal | 1,000 | 0.982 | 0.993 | 31.572 | 32.552 |
| 4 | ICA-FIX | 100 | 0.768 | 0.945 | 22.286 | 21.329 |
| 5 | ICA-FIX | 400 | 0.970 | 0.987 | 27.620 | 30.049 |
| 6 | ICA-FIX | 1,000 | 0.991 | 0.997 | 31.722 | 34.709 |
| Aggregation: *Mean* |  |  |  |  |  |  |
| 7 | Minimal | 100 | 0.954 | 0.954 | 19.587 | 19.587 |
| 8 | Minimal | 400 | 0.987 | 0.987 | 27.963 | 27.963 |
| 9 | Minimal | 1,000 | 0.993 | 0.993 | 32.530 | 32.530 |
| 10 | ICA-FIX | 100 | 0.95 | 0.95 | 21.301 | 21.301 |
| 11 | ICA-FIX | 400 | 0.987 | 0.987 | 30.044 | 30.044 |
| 12 | ICA-FIX | 1,000 | 0.997 | 0.997 | 34.768 | 34.768 |

The parcellation granularity was an important factor for the identification performance. In all cases, ***I_acc_*** increased with increasing parcellation granularity. ***I_acc_*** values were less influenced by parcellation granularity for *Mean* aggregation, ranging from 0.954 to 0.997 for both data quality configurations (Table 2). However, for *EV* aggregation, the parcellation granularity had a more pronounced impact in voxel-wise denoising for both data quality configurations (*Minimal*: 0.802-0.982; *ICA-FIX*: 0.768-0.991). For the parcellation granularity of 400 regions, all identification accuracies were above .94, increasing to 0.982 for 1000 regions. The parcellation granularity effect was also prominent in ***I_diff_***(Table 2). The performance obtained with the *ICA-FIX* data configuration was better compared with the one obtained from the *Minimal* data configuration, except for the *EV* aggregation in voxel-wise denoising, where the obtained performance was similar for both data quality configurations.

As expected, the denoising level did not influence ***I_acc_***or ***I_diff_*** for *Mean* aggregation, since the same performance was achieved between the two denoising levels in each parcellation granularity and each data configuration. The performance of the two aggregation methods (*Mean* and *EV*) was found to be almost equal when denoising at the regional level when the same data configuration and parcellation granularity were used.

### Predictiveness of rs-FC denoised at the voxel- or region-level

In the prediction analysis, age showed the strongest average correlation (0.286) between predicted and actual scores across all conditions (Figure 2). Among the cognitive scores, fluid intelligence had the strongest correlation (0.199), while Working Memory and Reading obtained 0.127 and 0.119, respectively.

**Figure 2.**
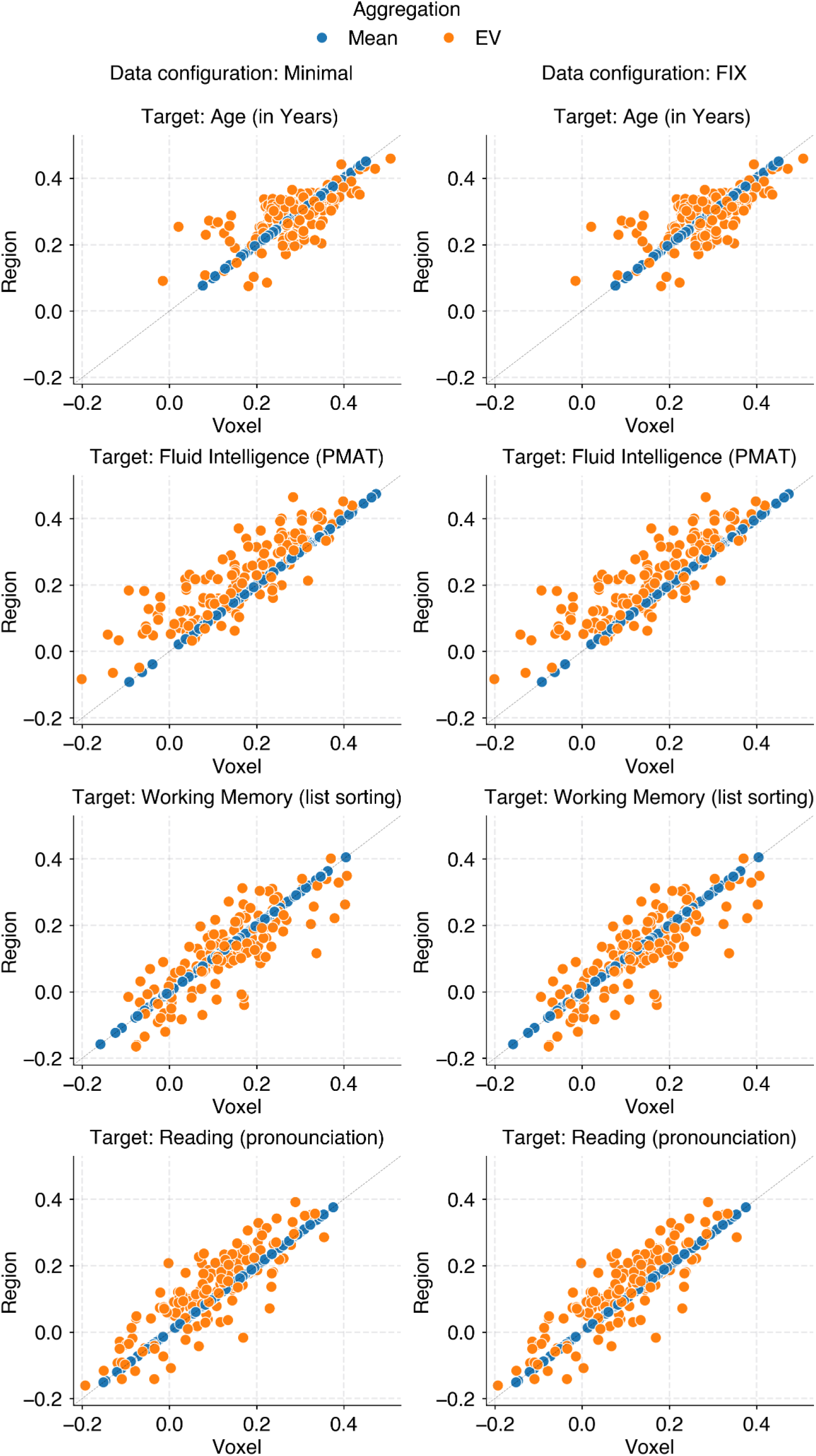
Predictability based on rs-FC denoised voxel-wise or region-wise in both data quality configurations. Pearson’s r between the predicted and observed values of each target, generated over 5-fold x 5-repeats in each of the three parcellation granularities are shown. Rs-FC was averaged across both runs (LR and RL) and both sessions (R1 and R2) per subject. Mean aggregation (blue) yielded identical predictions between the two denoising levels for all targets, as illustrated by their position on the diagonal. EV aggregation (orange) yielded slightly higher predictions for region-wise denoising for fluid intelligence and reading.

Between the two aggregation metrics, small differences in average correlation between predicted and actual scores were also observable for the targets fluid intelligence (Mean: 0.215 / EV: 0.183) and Reading (Mean: 0.128 / EV: 0.111). As expected, there were no differences between voxel- and region-level denoising for *Mean* aggregation (Figure 2). Regarding the *EV* aggregation method, there was no clear difference between voxel-level and region-level denoising for predicting age and working memory (Figure 2). However, region-level denoising showed a slight advantage for predicting fluid intelligence and reading ability (Figure 2).

Similar to the identification analyses, prediction performance increased with increasing parcellation granularity but only for the target *fluid intelligence* (Figure S3). This trend was most noticeable with *EV* aggregation, where voxel-level denoising showed increasing performance with increasing parcellation granularity. Region-level denoising increased only from 100 to 400 regions which was also seen in *Mean* aggregation performance.

### Assessment of motion in rs-FC denoised at the voxel- or region-level

As motion influences can still be present in preprocessed rs-FC (Siegel et al., 2017; Waller et al., 2017), we analyzed the relationship between typicality of the rs-FC and mean FD for each run. To this end, we sought to determine the association (Spearman correlation) between TFC and mean FD values (Figure 3). In this context, the difference between the data quality configurations was of particular interest, as they include different degrees of motion correction.

**Figure 3.**
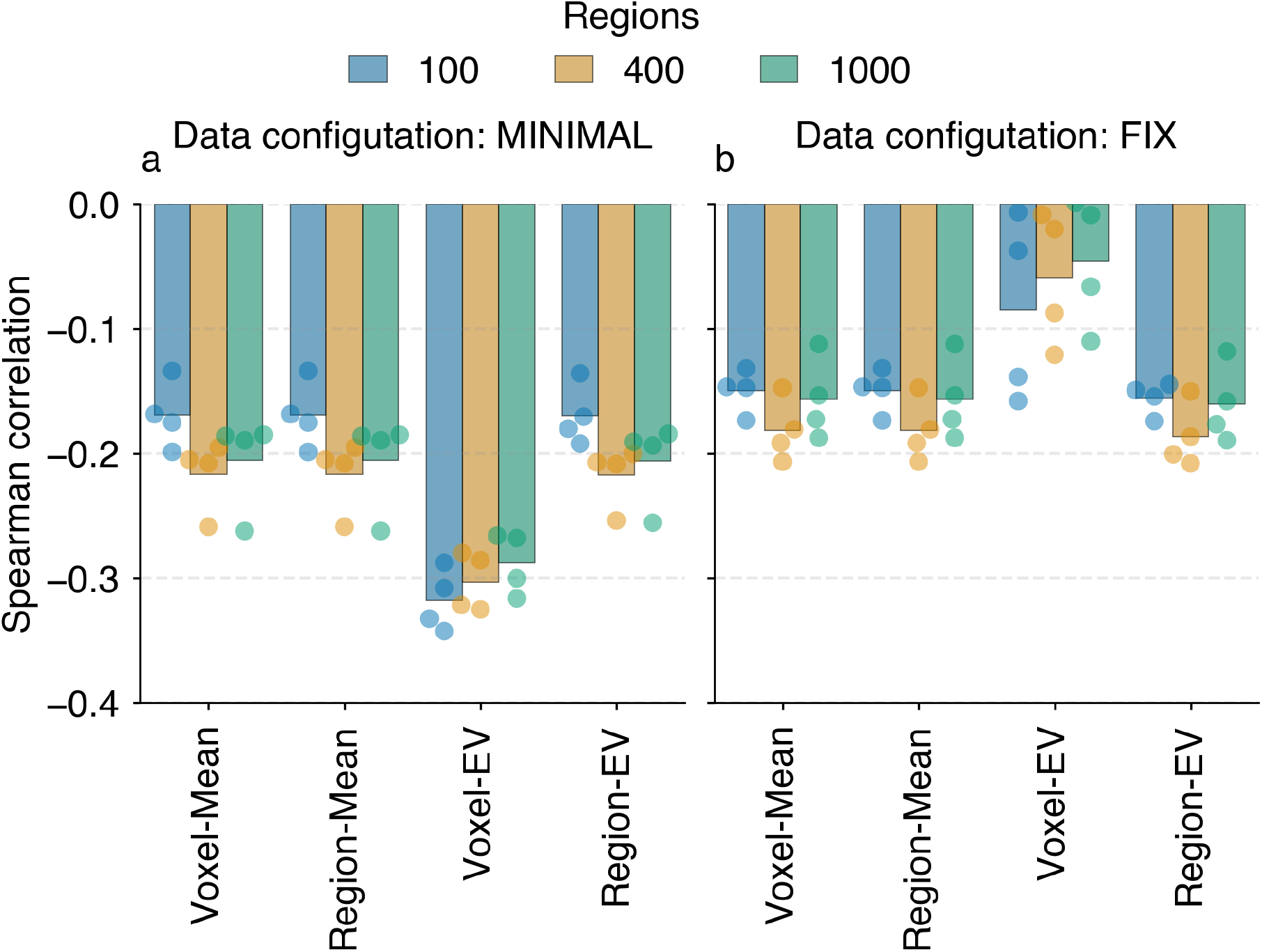
The association between TFC and mean FD values for all four phase encoding runs in both the Minimal data configuration (A) and ICA-FIX dataconfiguration (B) is presented across parcellation granularities. Each bar represents the average of all data points. (A) The association was found to be stable for both Mean aggregation and EV aggregation denoised at the region level. However, the association with motion was increased in the EV aggregated rs-FC denoised at the voxel level. (B) The values in the ICA-FIX dataset for Mean aggregation and EV aggregation denoised at the region level were stable and exhibited a slight decrease in comparison to those of the Minimal data configuration. The EV aggregated rs-FC denoised at the voxel level demonstrated the lowest overall association with motion.

As expected, for *Mean* aggregation, the same association values were obtained for both voxel- and region-level denoising ranging between -0.112 to -0.206 (*ICA-FIX*) and -0.133 to -0.262 (*Minimal*). For the *EV* aggregation, rs-FC was also more strongly associated with motion in the *Minimal* data configuration (-0.135 to -0.342), compared to the association obtained in the *ICA-FIX* data configuration (0.001 to -0.208). Importantly, in all the cases, the motion correlation was lower when using the *ICA-FIX* data configuration (0.001 to -0.206) compared to the *Minimal* data configuration (-0.133 to -0.342). Interestingly, this difference in data quality configurations was largest in the voxel-level denoised rs-FC using *EV* aggregation.

### Comparison of pipeline runtimes

The comparison of time required to preprocess and calculate the rs-FC matrices showed that using voxel-level denoising led to a 1.3-fold increase compared to region-level denoising, irrespective of the aggregation method or parcellation granularity (Figure 4). When comparing the two aggregation methods, using EV aggregation generally led to a 1.7-fold increase in computation time compared to *Mean* aggregation. An influence of parcellation granularity was observed exclusively in the context of *EV* aggregation. In this case, the usage of 100 regions demanded 1.7 times the computing time required by 1000 regions. Conversely, 1000 regions exhibited a computing time that was nearly comparable to that required for *Mean* aggregation.

**Figure 4.**
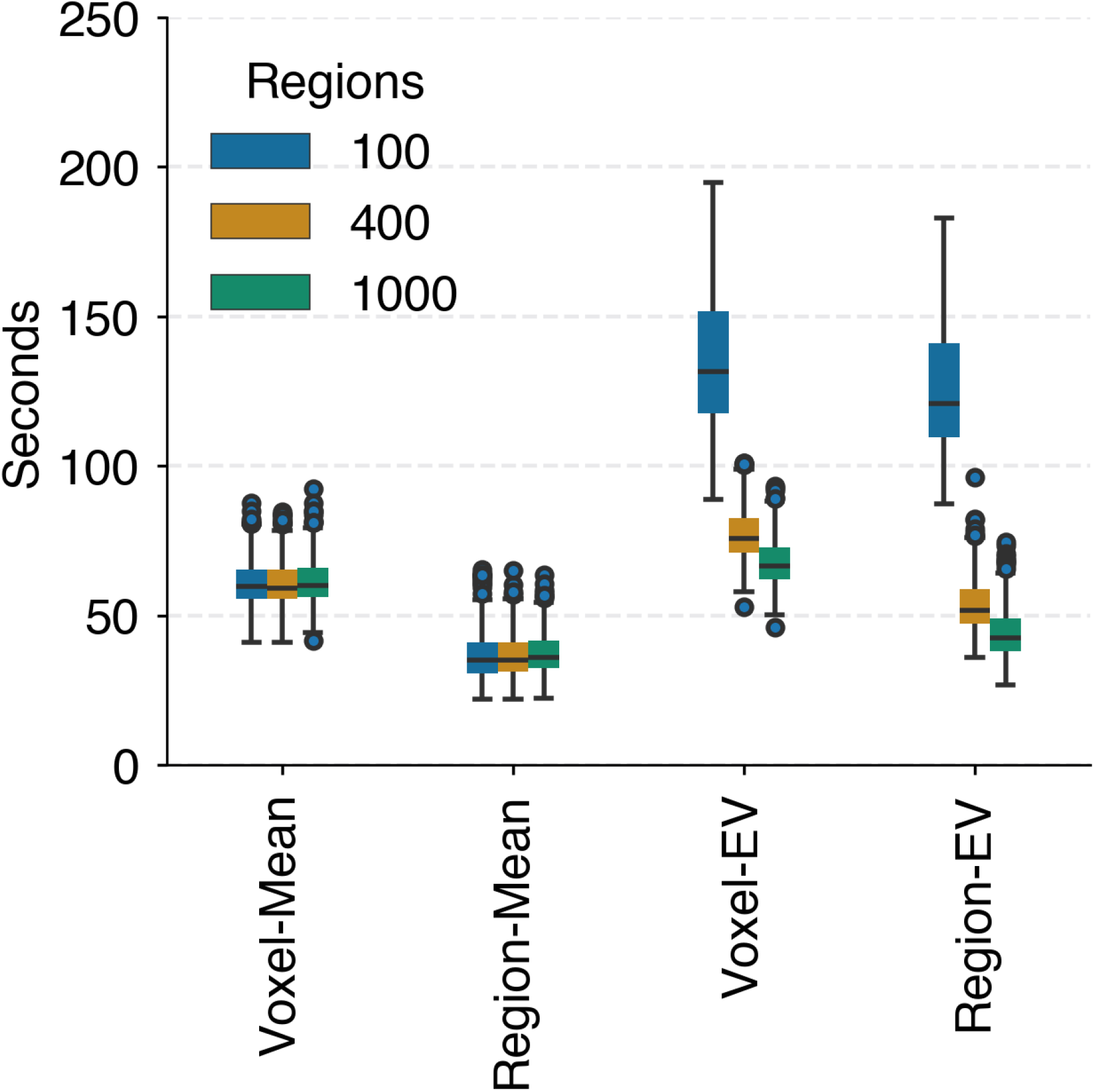
Pipeline runtime per subject (n=370) shows a decreasing trend with finer granularities in EV aggregation. The region-wise denoising pipeline was faster than the respective voxel-wise pipeline in all cases.

## Discussion

Here, our main goal was to systematically compare denoising on the aggregated region-level versus voxel-level BOLD time series in resting-state processing pipelines for individual-level rs-FC. To take into account other factors influencing rs-FC, we additionally considered two aggregation strategies to obtain regional time series (*Mean* and *EV*), three levels of cortical parcellation granularity (*100*, *400*, and *1,000* regions), and two data quality configurations (*Minimal* and *ICA-FIX*). Individual differences in the resulting rs-FC were examined using data from 370 unrelated subjects from the HCP-YA dataset. To assess the utility of the ensuing rs-FC, we evaluated two aspects of individual-level rs-FC; (1) individual specificity quantifying the stability of rs-FC across scans, and (2) the predictability of age, fluid intelligence, working memory, and reading ability. Additionally, we assessed the rs-FCs association with motion as well as computational resources required per pipeline execution. Our results indicated equal or higher performance for region-level denoising with more stable performance and reduced runtime in the Mean aggregation strategy. The general performance, especially in individual specificity increased with increasing parcellation granularity. Not surprisingly, rs-FC from the *Mean* aggregation yields equivalent results for both denoising levels across analyses of individual differences. For rs-FC from *EV* aggregation, the results between the two denoising levels were dissimilar. This dissimilarity was modulated by the parcellation granularity.

### Individual specificity of rs-FC denoised at voxel- or regional level

The useability of whole-brain rs-FC depends on capturing stable individual aspects of brain connectivity which allow reliable identification of the individual across multiple scans (Finn & Rosenberg, 2021). Overall the two individual specificity metrics, identification accuracy and differential identifiability, were within the ranges of previous reports (Amico & Goñi, 2018; Finn et al., 2015; Sasse et al., 2023). In particularly, the high identification accuracy scores (0.95-0.99; see Table 2) are to be expected given the high data quality and long scan duration of the HCP-YA dataset (Airan et al., 2016; Horien et al., 2018). As expected, the performance of *Mean* aggregation for *voxel*- and *region*-level denoising was equal in all cases. As pointed out by Friston et al. (2006), from the mathematical point of view, this is expected, as long as all steps performed are linear transformations of the data.

The individual specificity improved with finer parcellation granularity. This is in line with previous suggestions that a finer parcellation granularity benefits the identification of individuals in rs-FC (Airan et al., 2016; Finn et al., 2015). Especially, the finer parcellations (*400* and *1000* regions) led to almost similar individual specificity across denoising levels and aggregation strategies. As finer granularities contain smaller regions, they are more likely to enclose functionally homogeneous voxels. In cases like this, the behavior of *Mean* and *EV* aggregation has been suggested to be similar (Craddock et al., 2012; Saxe et al., 2006).

However, the coarser parcellation (*100* regions) showed a noteworthy difference between *voxel*- and *region*-level denoising for *EV* aggregation. This difference was characterized by reduced ***I_acc_***and increased ***I_diff_*** in *voxel*-level denoised rs-FC. Although most reports on individual specificity focus on *Mean* aggregated rs-FC, Airan et al.’s (2016) comparison of *Mean* and *EV* aggregation reported no noteworthy differences between the two aggregation strategies across a similar range of parcellation granularities. Their investigation focused on the amount of data (acquisition minutes) needed to maximize individual specificity in a sample of 23 subjects. To this end, they compared the two aggregation strategies in a 1,000-region Craddock parcellation (Craddock et al., 2012). Their finding of similarity between Mean and EV is consistent with our findings of individual differences at the finest parcellation granularity (1,000 regions). In addition, we have shown that the difference between the two aggregation strategies is greater at coarse parcellation granularity and when using a larger sample size with a similar amount of data per subject. In particular, the sample size has been suggested to affect measures of individual specificity, as a larger number of subjects also increase the likelihood of observing similar rs-FC between subjects (Li et al., 2021; Waller et al., 2017).

Borrowing from the previous arguments of Craddock et al. (2012) and Saxe et al. (2006), the *EV* will select a subset of voxel times series per region in heterogeneous regions, which may lead to the selection of different voxels across multiple scans. This session specificity could lead to an increased likelihood that single individuals get misidentified in a binary test-retest metric such as ***I_acc_***, where a single mismatch is enough to influence the overall score. The differential identifiability’s ability to quantify the individual specificity on a continuous scale across the whole dataset could still be rather high in this scenario, as long as the mismatch is caused by only one or a few other subjects being more similar.

The suggested behavior of the *EV* only led to individual differences in voxel-level denoising. Aggregating the voxel time series first and then denoising the *region*-level time series led to individual specificity performance similar to *Mean* aggregation. Since denoising involves detrending, effectively demeaning the time series, the *EV* is influenced to select slightly different variance components depending on the denoising level. In the case of *region*-level denoising, aggregating the voxel time series might lead to the EV decomposition selecting the regional mean signal, since this component explains the largest amount of variance.

### Predictability from rs-FC denoised at voxel- or regional level

Another aspect of rs-FC for effective use in clinical applications is its utility in predicting individual-level demographics and phenotypes (Finn & Rosenberg, 2021). In our study, the prediction accuracies (Pearson’s r) between 0.286 and 0.119 (Figure 2) were in line with previously reported ranges for whole-brain rs-FC-based predictions (He et al., 2020; Sasse et al., 2023). The similarity observed in individual specificity between *voxel*- and *region*-level denoising when using Mean aggregation was also observed for the prediction accuracies across all targets. Thus, the predictive utility of rs-FC did not change irrespective of voxel- or region-level denoising. Nevertheless, *voxel*-level denoising is widely used in brain and behavioural studies as the common toolboxes SPM, FSL, CONN use it as their default. However, due to the mathematical equality, there seems to be no obvious reason to choose one or the other in terms of prediction performance.

The differences observed in the *EV* aggregation were present only in the targets fluid intelligence and reading ability. The rs-FC that is optimal for individual specificity is not necessarily optimal for prediction accuracy because different aspects of the signal might contribute to these different goals (Finn & Constable, 2016; Noble et al., 2019). Similar or slightly lower prediction accuracies were obtained for *EV-aggregated* rs-FC denoised at the *voxel*-level. This suggests that *EV-aggregated* rs-FC does not offer an advantage in prediction analysis. Thus, the potentially interesting ability to capture session-specific details by *EV* aggregation did not uncover patterns that would lead to better predictions of the behavioral targets or age.

Of the four targets, only fluid intelligence was affected by the parcellation granularity. Previous studies have demonstrated that the frontal-parietal network is a driving factor for individual specificity as well as fluid intelligence (Amico & Goñi, 2018; Finn et al., 2015). Our results suggest that this alignment between individual specificity and predictability is only meaningful for the prediction of fluid intelligence scores while other targets might rest on different aspects of the rs-FC. In this target, the lowest parcellation granularity (*100* regions) performed worst across the denoising levels and aggregation strategies. The slight improvements with finer granularities suggest that the relevant signal might not be adequately captured at coarse granularities. Interestingly, increasing to finer parcellation did not lead to improved performance at *400 and 1,000* regions. Here, it is possible that in large feature spaces, like the *1,000*-region parcellation, the signal might become diluted, i.e., many features might not be associated with the target, and consideration of the curse-of-dimensionality (Hastie et al., 2009) can harm predictive performance (Schulz et al., 2024).

### Additional considerations

#### Motion association

Previous studies suggest that residual motion may influence the individual specificity and predictability of rs-FC (Siegel et al., 2017; Waller et al., 2017). However, others suggest that this influence is likely to be minimized with a sufficient amount of data per subject (Airan et al., 2016; Horien et al., 2018). Our main findings support this view, as we observed stable individual specificity and predictive performance regardless of the degree of preprocessing (*Minimal* or *ICA-FIX*) applied to the raw data by the HCP (Glasser et al., 2013; Smith et al., 2013). In addition to our main findings, we examined the quality of each of the rs-FC runs and its relation to motion by associating the typicality of the rs-FC of each run with the respective average FD (Kopal et al., 2020). Some stable and low degree of association remained in all *Mean* aggregation rs-FCs and the *EV* aggregated rs-FC using *region*-level denoising. These associations have also been reported previously (Kopal et al., 2020). Interestingly, the *voxel*-level denoising in *EV* aggregated rs-FC displayed differing patterns between the two data quality configurations. In the *Minimal* data configuration, the association to motion was the highest, while the associations in the *ICA-FIX* data configuration were close to zero. This effect was stable across the parcellation granularities (see Figure 3). We take this result as an indicator that, indeed, the quality of the *ICA-FIX* data configuration is less motion-associated. However, this effect does not translate into the performance in our prediction analysis.

#### Pipeline Runtime

In general, the denoising of the regional time series requires less computational time, as shown by the runtime of the different pipelines. The most noticeable difference in runtime was between the aggregation strategies. The *Mean* aggregation strategy had a constant runtime across all levels of parcellation granularity. In contrast, the *EV* aggregation took more time when coarser parcellation granularities (i.e. more voxels per region) were involved. This difference is likely due to the *Mean* strategy being easier to perform, making it efficient regardless of the number of voxels in a region. In contrast, singular value decomposition determines all eigenvectors, of which the first is projected back into the time series domain, which is much more time-consuming. This is especially true when a larger number of voxels are involved in the calculation, as the time complexity of standard EV implementations increases with the number of voxels.

### Limitations and considerations for future work

It should be noted that while we discuss the differences between voxel-wise and region-wise denoising, the ICA-FIX pipeline by the HCP was applied to the voxel time series. Hence, our results only account for the additional denoising steps of detrending, bandpass filtering, and linear nuisance regression. Future studies are needed to demonstrate that the obtained results are generalizable to other datasets with differing data quality and scan durations, which are known to influence individual difference analyses (Airan et al., 2016; Horien et al., 2018). Since *Mean* aggregation should be the same for voxel- and region-level denoising based on firm mathematical reasons and supported by our empirical demonstration, it should lead to similar outcomes in other datasets. However, the performance of the *EV* might vary depending on multiple factors such as motion content and choice of region size. The influence of these factors were already observable in the current dataset. Note that while ICA-FIX leads to clean data and is a HCP-specific configuration, the results from the minimally processed data configuration might be considered more similar to standard quality clinical datasets. Here we employed parcellation schemes with three levels of parcellation granularity and found that the largest improvement in identification performance happened between 100 and 400 regions. It might be of interest for future investigations to probe the space between these two granularities. Lastly, as a group-level parcellation was used, it is possible that voxel-to-region assignment across subjects was suboptimal which might have influenced the *EV* aggregation more than the *Mean* aggregation. Individual-specific region definition approaches might provide better region specification (Kong et al., 2021). Furthermore, additional steps can be performed in rs-FC processing (e.g., scrubbing) that we did not consider. Future work might consider those aspects more closely.

## Conclusion

Based on our findings, we encourage the potential of denoising time series at the regional level for the analysis of individual differences. Particularly for *Mean* aggregation, as it provides equivalent results and is computationally more efficient. This offers substantial benefits in computational runtime without compromising the individual specificity and predictability of *Mean* aggregated rs-FC. While *EV* aggregation might be an interesting alternative, the sensitivity to residual motion should be closely monitored and evaluated, especially with voxel-wise denoising. Lastly, the choice of parcellation granularity (i.e., region size) should be carefully considered specifically for *EV* aggregation, as individual differences may suffer in coarser parcellations. With this work, we hope to provide evidence to guide researchers in choosing the most appropriate and efficient steps for investigating brain-behavior associations in future investigations.

## Acknowledgments

This work was partly supported by the Helmholtz Portfolio Theme “Supercomputing and Modelling for the Human Brain”. Data were provided [in part] by the Human Connectome Project, WU-Minn Consortium (Principal Investigators: David Van Essen and Kamil Ugurbil; 1U54MH091657) funded by the 16 NIH Institutes and Centers that support the NIH Blueprint for Neuroscience Research; and by the McDonnell Center for Systems Neuroscience at Washington University.

## Author contributions

**Tobias Muganga**: Conceptualization, Data Curation, Formal Analysis, Investigation, Methodology, Project Administration, Software, Validation, Visualization, Writing - Original Draft Preparation, Writing - Review & Editing

**Leonard Sasse**: Data Curation, Formal Analysis, Methodology, Software, Validation, Visualization, Writing - Review & Editing

**Daouia I. Larabi**: Conceptualization, Supervision, Writing - Original Draft Preparation, Writing - Review & Editing

**Nicolás Nieto**: Visualization, Writing - Review & Editing

**Julian Caspers**: Supervision, Writing - Review & Editing

**Simon B. Eickhoff**: Conceptualization, Funding Acquisition, Supervision, Writing - Review & Editing

**Kaustubh R. Patil**: Conceptualization, Funding Acquisition, Investigation, Methodology, Supervision, Validation, Writing - Original Draft Preparation, Writing - Review & Editing

## Supplementary Material

**Figure S1.**
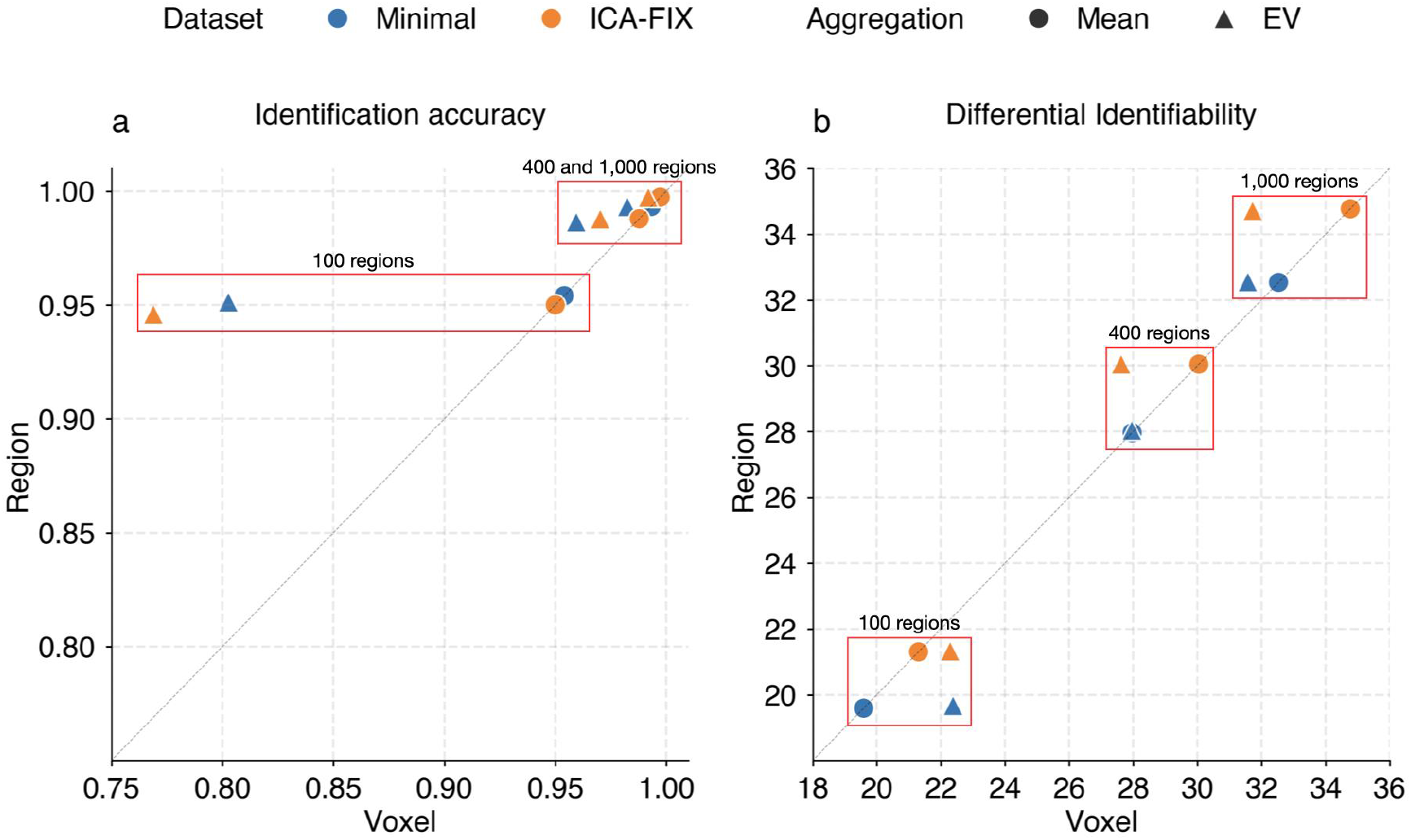
Comparing identification performance of rs-FC denoised at the voxel- versus region-level across Mean or EV aggregation (shape) and three parcellation granularities in both data quality configurations (color). (a) Identification accuracy increases with finer granularities. Differences between the denoising levels are observed only in the case of EV aggregation. (b): Differential Identifiability also increases with finer parcellation. Deviations from the diagonal only occur in EV aggregation in favor of voxel-level denoising at 100 regions and equal or better for region-level denoising at 400 and 1,000 regions.

**Figure S2.**
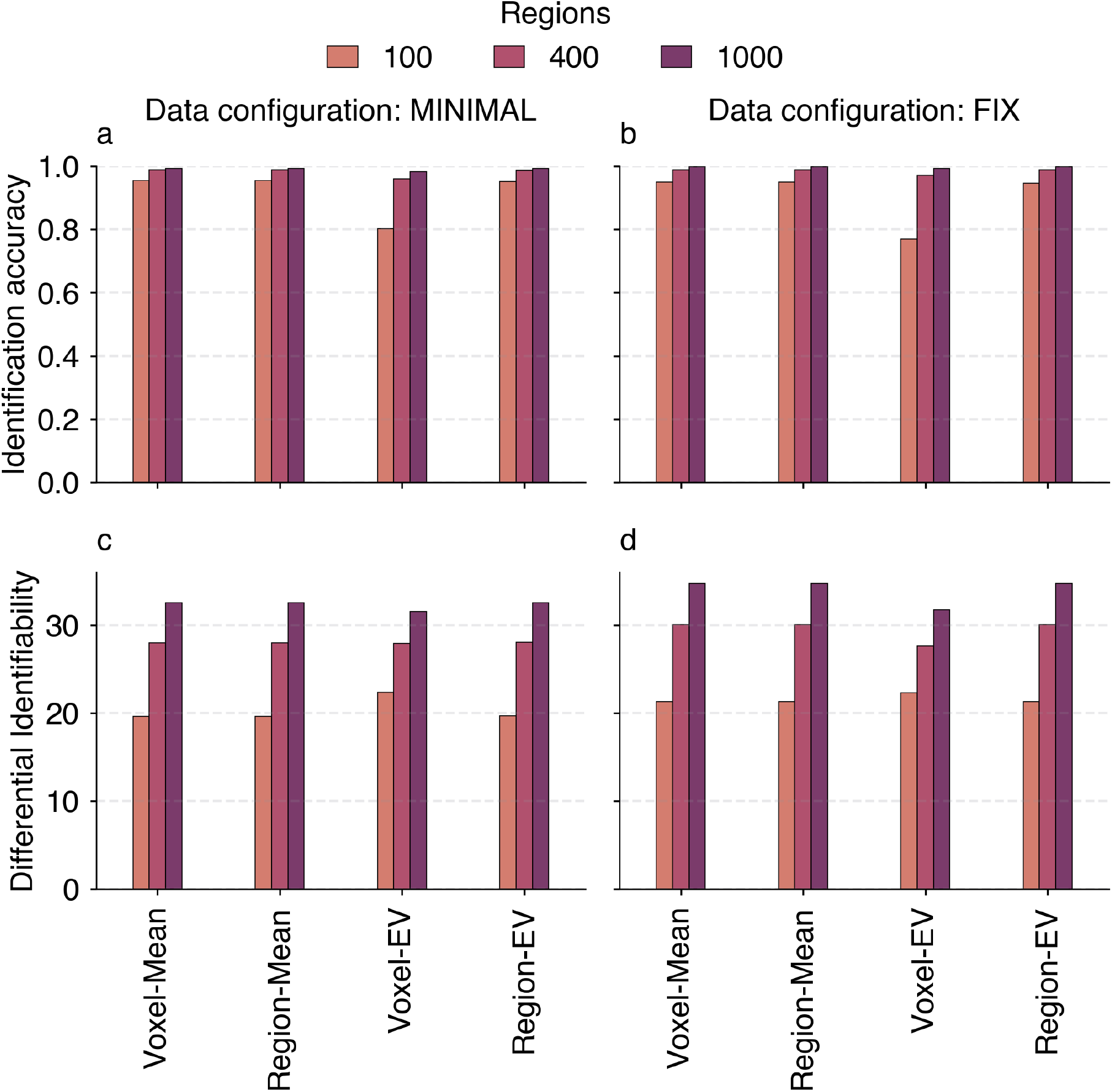
Overview of individual specificity of rs-FC across parcellation granularities. Results are shown for rs-FC generated from Minimal (a & c) and ICA-FIX (b & d) data configuration.

**Figure S3.**
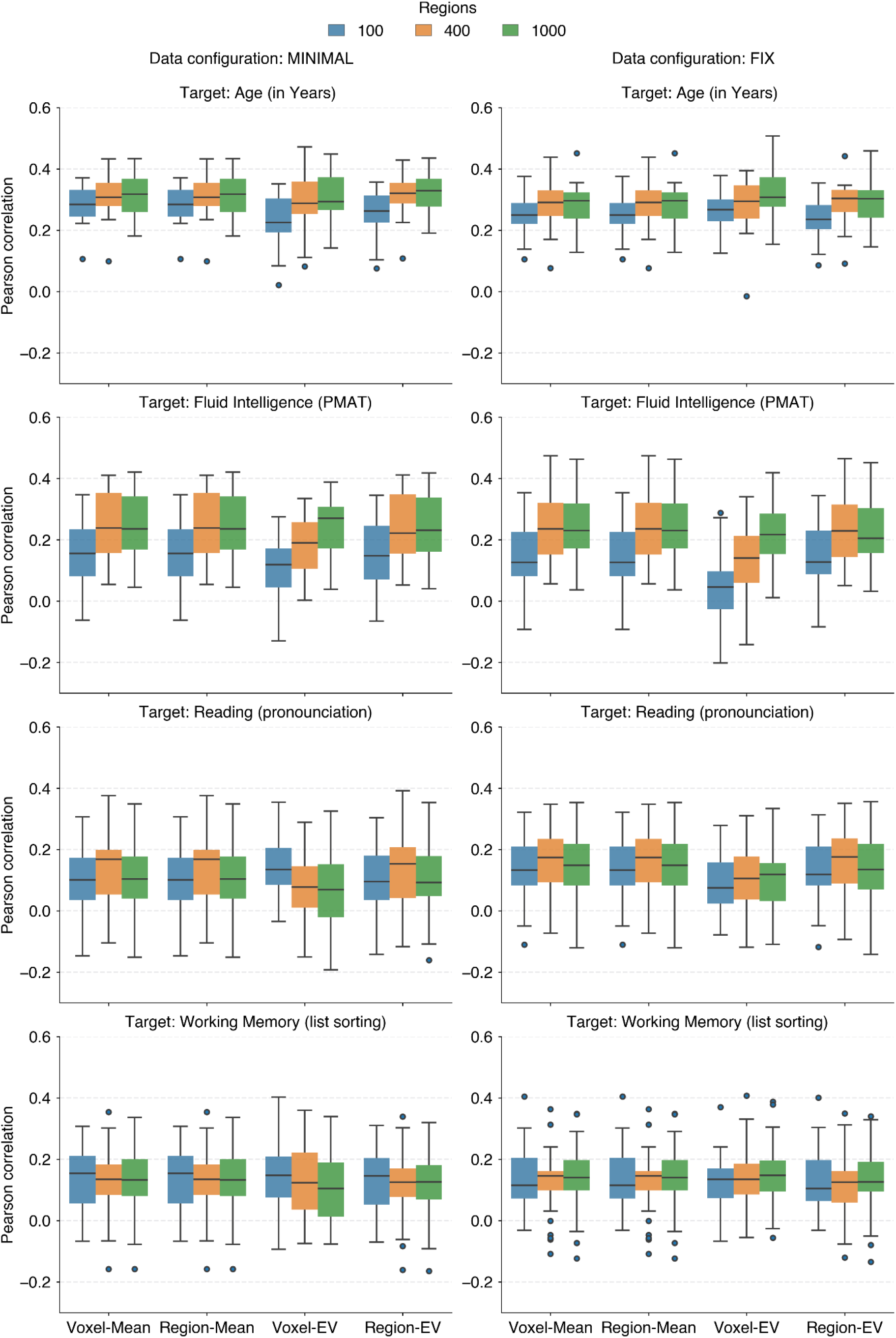
Overview of predictability based on rs-FC denoised voxel-wise or region-wise in both data quality configurations. Pearson’s r between the predicted and observed values of each target, generated over 5-fold x 5-repeats in each of the three parcellation granularities are shown. Rs-FC was averaged across both runs (LR and RL) and both sessions (R1 and R2) per subject.

## References

1. Abraham, A., Pedregosa, F., Eickenberg, M., Gervais, P., Mueller, A., Kossaifi, J., Gramfort, A., Thirion, B., & Varoquaux, G. (2014). Machine learning for neuroimaging with scikit-learn. Frontiers in Neuroinformatics, 8. 10.3389/fninf.2014.00014

2. Airan, R. D., Vogelstein, J. T., Pillai, J. J., Caffo, B., Pekar, J. J., & Sair, H. I. (2016). Factors affecting characterization and localization of interindividual differences in functional connectivity using MRI. Human Brain Mapping, 37(5), 1986. 10.1002/hbm.23150

3. Amico, E., & Goñi, J. (2018). The quest for identifiability in human functional connectomes. Scientific Reports 2018 *8*:1, *8*(1), 1–14. 10.1038/s41598-018-25089-1

4. Behzadi, Y., Restom, K., Liau, J., & Liu, T. T. (2007). A component based noise correction method (CompCor) for BOLD and perfusion based fMRI. NeuroImage, 37(1), 90–101. 10.1016/j.neuroimage.2007.04.042

5. Bianciardi, M., Fukunaga, M., van Gelderen, P., Horovitz, S. G., de Zwart, J. A., Shmueli, K., & Duyn, J. H. (2009). Sources of fMRI signal fluctuations in the human brain at rest: A 7T study. Magnetic Resonance Imaging, 27(8), 1019. 10.1016/J.MRI.2009.02.004

6. Biswal, B. B., Mennes, M., Zuo, X. N., Gohel, S., Kelly, C., Smith, S. M., Beckmann, C. F., Adelstein, J. S., Buckner, R. L., Colcombe, S., Dogonowski, A. M., Ernst, M., Fair, D., Hampson, M., Hoptman, M. J., Hyde, J. S., Kiviniemi, V. J., Kötter, R., Li, S. J., … Milham, M. P. (2010). Toward discovery science of human brain function. Proceedings of the National Academy of Sciences of the United States of America, 107(10), 4734–4739. 10.1073/PNAS.0911855107/SUPPL_FILE/PNAS.200911855SI.PDF

7. Biswal, B., Yetkin, F. Z., Haughton, V. M., & Hyde, J. S. (1995). Functional connectivity in the motor cortex of resting human brain using echo-planar mri. Magnetic Resonance in Medicine, 34(4), 537–541. 10.1002/MRM.1910340409

8. Bright, M. G., Tench, C. R., & Murphy, K. (2017). Potential pitfalls when denoising resting state fMRI data using nuisance regression. NeuroImage, 154, 159–168. 10.1016/J.NEUROIMAGE.2016.12.027

9. Ciric, R., Wolf, D. H., Power, J. D., Roalf, D. R., Baum, G. L., Ruparel, K., Shinohara, R. T., Elliott, M. A., Eickhoff, S. B., Davatzikos, C., Gur, R. C., Gur, R. E., Bassett, D. S., & Satterthwaite, T. D. (2017). Benchmarking of participant-level confound regression strategies for the control of motion artifact in studies of functional connectivity. NeuroImage, 154, 174–187. 10.1016/J.NEUROIMAGE.2017.03.020

10. Craddock, R. C., Holtzheimer, P. E., Hu, X. P., & Mayberg, H. S. (2012). A whole brain fMRI atlas generated via spatially constrained spectral clustering. Human Brain Mapping, 33(8), 1914–1928. 10.1002/HBM.21333

11. Demertzi, A., Kucyi, A., Ponce-Alvarez, A., Keliris, G. A., Whitfield-Gabrieli, S., & Deco, G. (2022). Functional network antagonism and consciousness. Network Neuroscience, 6(4), 998–1009. 10.1162/netn_a_00244

12. Dubois, J., & Adolphs, R. (2016). Building a Science of Individual Differences from fMRI. Trends in Cognitive Sciences, 20(6), 425–443. 10.1016/j.tics.2016.03.014

13. Eickhoff, S. B., Yeo, B. T. T., & Genon, S. (2018). Imaging-based parcellations of the human brain. Nature Reviews Neuroscience 2018 *19*:11, *19*(11), 672–686. 10.1038/s41583-018-0071-7

14. Finn, E. S., & Constable, R. T. (2016). Individual variation in functional brain connectivity: Implications for personalized approaches to psychiatric disease. Dialogues in Clinical Neuroscience, 18(3), 277–287. 10.31887/DCNS.2016.18.3/EFINN

15. Finn, E. S., & Rosenberg, M. D. (2021). Beyond fingerprinting: Choosing predictive connectomes over reliable connectomes. NeuroImage, 239. 10.1016/J.NEUROIMAGE.2021.118254

16. Finn, E. S., Shen, X., Scheinost, D., Rosenberg, M. D., Huang, J., Chun, M. M., Papademetris, X., & Constable, R. T. (2015). Functional connectome fingerprinting: Identifying individuals using patterns of brain connectivity. Nature Neuroscience 2015 *18*:11, *18*(11), 1664–1671. 10.1038/nn.4135

17. Fox, M. D., & Raichle, M. E. (2007). Spontaneous fluctuations in brain activity observed with functional magnetic resonance imaging. Nature Reviews Neuroscience 2007 8:9, 8(9), 700–711. 10.1038/nrn2201

18. Fox, M. D., Zhang, D., Snyder, A. Z., & Raichle, M. E. (2009). The Global Signal and Observed Anticorrelated Resting State Brain Networks. Journal of Neurophysiology, 101(6), 3270–3283. 10.1152/jn.90777.2008

19. Friston, K. J. (2011). Functional and Effective Connectivity: A Review. Brain Connectivity, 1(1), 13–36. 10.1089/brain.2011.0008

20. Friston, K. J., Rotshtein, P., Geng, J. J., Sterzer, P., & Henson, R. N. (2006). A critique of functional localisers. NeuroImage, 30(4), 1077–1087. 10.1016/j.neuroimage.2005.08.012

21. Friston, K. J., Williams, S., Howard, R., Frackowiak, R. S. J., & Turner, R. (1996). Movement-related effects in fMRI time-series. Magnetic Resonance in Medicine, 35(3), 346–355. 10.1002/MRM.1910350312

22. Geerligs, L., Rubinov, M., Tyler, L. K., Brayne, C., Bullmore, E. T., Calder, A. C., Cusack, R., Dalgleish, T., Duncan, J., Henson, R. N., Matthews, F. E., Marslen-Wilson, W. D., Rowe, J. B., Shafto, M. A., Campbell, K., Cheung, T., Davis, S., Geerligs, L., Kievit, R., … Henson, R. N. (2015). State and Trait Components of Functional Connectivity: Individual Differences Vary with Mental State. Journal of Neuroscience, 35(41), 13949–13961. 10.1523/JNEUROSCI.1324-15.2015

23. Glasser, M. F., Sotiropoulos, S. N., Wilson, J. A., Coalson, T. S., Fischl, B., Andersson, J. L., Xu, J., Jbabdi, S., Webster, M., Polimeni, J. R., Essen, D. C. V., & Jenkinson, M. (2013). The minimal preprocessing pipelines for the Human Connectome Project. NeuroImage, 80, 105–124. 10.1016/J.NEUROIMAGE.2013.04.127

24. Gordon, E. M., Laumann, T. O., Gilmore, A. W., Newbold, D. J., Greene, D. J., Berg, J. J., Ortega, M., Hoyt-Drazen, C., Gratton, C., Sun, H., Hampton, J. M., Coalson, R. S., Nguyen, A. L., McDermott, K. B., Shimony, J. S., Snyder, A. Z., Schlaggar, B. L., Petersen, S. E., Nelson, S. M., & Dosenbach, N. U. F. (2017). Precision Functional Mapping of Individual Human Brains. Neuron, 95(4), 791–807.e7. 10.1016/J.NEURON.2017.07.011

25. Grabner, G., Janke, A. L., Budge, M. M., Smith, D., Pruessner, J., & Collins, D. L. (2006). Symmetric atlasing and model based segmentation: An application to the hippocampus in older adults. Lecture Notes in Computer Science (Including Subseries Lecture Notes in Artificial Intelligence and Lecture Notes in Bioinformatics), 4191 LNCS-II, 58–66. 10.1007/11866763_8/COVER

26. Gratton, C., Laumann, T. O., Nielsen, A. N., Greene, D. J., Gordon, E. M., Gilmore, A. W., Nelson, S. M., Coalson, R. S., Snyder, A. Z., Schlaggar, B. L., Dosenbach, N. U. F., & Petersen, S. E. (2018). Functional Brain Networks Are Dominated by Stable Group and Individual Factors, Not Cognitive or Daily Variation. Neuron, 98(2), 439–452.e5. 10.1016/j.neuron.2018.03.035

27. Hamdan, S., More, S., Sasse, L., Komeyer, V., Patil, K. R., & Raimondo, F. (2023). *Julearn: An easy-to-use library for leakage-free evaluation and inspection of ML models* (arXiv:2310.12568). arXiv. 10.48550/arXiv.2310.12568

28. Hastie, T., Tibshirani, R., & Friedman, J. (2009). The Elements of Statistical Learning: Data Mining, Inference, and Prediction, Second Edition (Springer Series in Statistics).

29. He, T., Kong, R., Holmes, A. J., Nguyen, M., Sabuncu, M. R., Eickhoff, S. B., Bzdok, D., Feng, J., & Yeo, B. T. T. (2020). Deep neural networks and kernel regression achieve comparable accuracies for functional connectivity prediction of behavior and demographics. NeuroImage, 206, 116276. 10.1016/J.NEUROIMAGE.2019.116276

30. Horien, C., Noble, S., Finn, E. S., Shen, X., Scheinost, D., & Constable, R. T. (2018). Considering factors affecting the connectome-based identification process: Comment on Waller et al. NeuroImage, 169, 172–175. 10.1016/J.NEUROIMAGE.2017.12.045

31. Horien, C., Shen, X., Scheinost, D., & Constable, R. T. (2019). The individual functional connectome is unique and stable over months to years. NeuroImage, 189, 676–687. 10.1016/j.neuroimage.2019.02.002

32. Kong, R., Li, J., Orban, C., Sabuncu, M. R., Liu, H., Schaefer, A., Sun, N., Zuo, X. N., Holmes, A. J., Eickhoff, S. B., & Yeo, B. T. T. (2019). Spatial Topography of Individual-Specific Cortical Networks Predicts Human Cognition, Personality, and Emotion. *Cerebral Cortex (New York*, NY), 29(6), 2533. 10.1093/CERCOR/BHY123

33. Kong, R., Yang, Q., Gordon, E., Xue, A., Yan, X., Orban, C., Zuo, X. N., Spreng, N., Ge, T., Holmes, A., Eickhoff, S., & Yeo, B. T. T. (2021). Individual-Specific Areal-Level Parcellations Improve Functional Connectivity Prediction of Behavior. *Cerebral Cortex (New York*, NY), 31(10), 4477. 10.1093/CERCOR/BHAB101

34. Kopal, J., Pidnebesna, A., Tomeček, D., Tintěra, J., & Hlinka, J. (2020). Typicality of functional connectivity robustly captures motion artifacts in rs-fMRI across datasets, atlases, and preprocessing pipelines. Human Brain Mapping, 41(18), 5325–5340. 10.1002/hbm.25195

35. Korhonen, O., Saarimäki, H., Glerean, E., Sams, M., Saramäki, J., & Zuo, X. (2017). Consistency of Regions of Interest as nodes of fMRI functional brain networks. Network Neuroscience, 1(3), 254–274. 10.1162/NETN_A_00013

36. Leopold, D. A., Murayama, Y., & Logothetis, N. K. (2003). Very Slow Activity Fluctuations in Monkey Visual Cortex: Implications for Functional Brain Imaging. Cerebral Cortex, 13(4), 422–433. 10.1093/cercor/13.4.422

37. Li, J., Kong, R., Liégeois, R., Orban, C., Tan, Y., Sun, N., Holmes, A. J., Sabuncu, M. R., Ge, T., & Yeo, B. T. T. (2019). Global Signal Regression Strengthens Association between Resting-State Functional Connectivity and Behavior. NeuroImage, 196, 126. 10.1016/J.NEUROIMAGE.2019.04.016

38. Li, K., Wisner, K., & Atluri, G. (2021). Feature selection framework for functional connectome fingerprinting. Human Brain Mapping, 42(12), 3717–3732. 10.1002/hbm.25379

39. Lindquist, M. A., Geuter, S., Wager, T. D., & Caffo, B. S. (2019). Modular preprocessing pipelines can reintroduce artifacts into fMRI data. Human Brain Mapping, 40(8), 2358. 10.1002/HBM.24528

40. Logothetis, N. K. (2003). The Underpinnings of the BOLD Functional Magnetic Resonance Imaging Signal. Journal of Neuroscience, 23(10), 3963–3971. 10.1523/JNEUROSCI.23-10-03963.2003

41. Macey, P. M., Macey, K. E., Kumar, R., & Harper, R. M. (2004). A method for removal of global effects from fMRI time series. NeuroImage, 22(1), 360–366. 10.1016/j.neuroimage.2003.12.042

42. Mantwill, M., Gell, M., Krohn, S., & Finke, C. (2022). Brain connectivity fingerprinting and behavioural prediction rest on distinct functional systems of the human connectome. Communications Biology, 5(1), 1–10. 10.1038/s42003-022-03185-3

43. More, S., Antonopoulos, G., Hoffstaedter, F., Caspers, J., Eickhoff, S. B., & Patil, K. R. (2023). Brain-age prediction: A systematic comparison of machine learning workflows. NeuroImage, 270, 119947. 10.1016/j.neuroimage.2023.119947

44. Noble, S., Scheinost, D., & Constable, R. T. (2019). A decade of test-retest reliability of functional connectivity: A systematic review and meta-analysis. NeuroImage, 203, 116157. 10.1016/J.NEUROIMAGE.2019.116157

45. Parkes, L., Fulcher, B., Yücel, M., & Fornito, A. (2018). An evaluation of the efficacy, reliability, and sensitivity of motion correction strategies for resting-state functional MRI. NeuroImage, 171. 10.1016/j.neuroimage.2017.12.073

46. Pedregosa, F., Varoquaux, G., Gramfort, A., Michel, V., Thirion, B., Grisel, O., Blondel, M., Prettenhofer, P., Weiss, R., Dubourg, V., Vanderplas, J., Passos, A., Cournapeau, D., Brucher, M., Perrot, M., & Duchesnay, E. (2011). Scikit-learn: Machine Learning in Python. Journal of Machine Learning Research, 12, 2825–2830.

47. Pernet, C. R., Wilcox, R., & Rousselet, G. A. (2013). Robust correlation analyses: False positive and power validation using a new open source matlab toolbox. Frontiers in Psychology, 3(JAN), 606. 10.3389/FPSYG.2012.00606/BIBTEX

48. Power, J. D., Plitt, M., Laumann, T. O., & Martin, A. (2017). Sources and implications of whole-brain fMRI signals in humans. NeuroImage, 146, 609–625. 10.1016/j.neuroimage.2016.09.038

49. Power, J. D., Schlaggar, B. L., & Petersen, S. E. (2014). Studying Brain Organization via Spontaneous fMRI Signal. Neuron, 84(4), 681–696. 10.1016/J.NEURON.2014.09.007

50. Power, J. D., Schlaggar, B. L., & Petersen, S. E. (2015). Recent progress and outstanding issues in motion correction in resting state fMRI. NeuroImage, 105, 536–551. 10.1016/J.NEUROIMAGE.2014.10.044

51. Salimi-Khorshidi, G., Douaud, G., Beckmann, C. F., Glasser, M. F., Griffanti, L., & Smith, S. M. (2014). Automatic denoising of functional MRI data: Combining independent component analysis and hierarchical fusion of classifiers. NeuroImage, 90, 449–468. 10.1016/j.neuroimage.2013.11.046

52. Sasse, L., Larabi, D. I., Omidvarnia, A., Jung, K., Hoffstaedter, F., Jocham, G., Eickhoff, S. B., & Patil, K. R. (2023). Intermediately synchronised brain states optimise trade-off between subject specificity and predictive capacity. Communications Biology, 6(1), 1–14. 10.1038/s42003-023-05073-w

53. Satterthwaite, T. D., Elliott, M. A., Gerraty, R. T., Ruparel, K., Loughead, J., Calkins, M. E., Eickhoff, S. B., Hakonarson, H., Gur, R. C., Gur, R. E., & Wolf, D. H. (2013). An improved framework for confound regression and filtering for control of motion artifact in the preprocessing of resting-state functional connectivity data. NeuroImage, 64, 240–256. 10.1016/j.neuroimage.2012.08.052

54. Saxe, R., Brett, M., & Kanwisher, N. (2006). Divide and conquer: A defense of functional localizers. NeuroImage, 30(4), 1088–1096. 10.1016/j.neuroimage.2005.12.062

55. Schaefer, A., Kong, R., Gordon, E. M., Laumann, T. O., Zuo, X.-N., Holmes, A. J., Eickhoff, S. B., & Yeo, B. T. T. (2018). Local-Global Parcellation of the Human Cerebral Cortex from Intrinsic Functional Connectivity MRI. Cerebral Cortex, 28(9), 3095–3114. 10.1093/CERCOR/BHX179

56. Schulz, M.-A., Bzdok, D., Haufe, S., Haynes, J.-D., & Ritter, K. (2024). Performance reserves in brain-imaging-based phenotype prediction. Cell Reports, 43(1). 10.1016/j.celrep.2023.113597

57. Shen, X., Tokoglu, F., Papademetris, X., & Constable, R. T. (2013). Groupwise whole-brain parcellation from resting-state fMRI data for network node identification. NeuroImage, 82, 403–415. 10.1016/j.neuroimage.2013.05.081

58. Siegel, J. S., Mitra, A., Laumann, T. O., Seitzman, B. A., Raichle, M., Corbetta, M., & Snyder, A. Z. (2017). Data Quality Influences Observed Links Between Functional Connectivity and Behavior. Cerebral Cortex, 27(9), 4492–4502. 10.1093/CERCOR/BHW253

59. Smith, S. M., Beckmann, C. F., Andersson, J., Auerbach, E. J., Bijsterbosch, J., Douaud, G., Duff, E., Feinberg, D. A., Griffanti, L., Harms, M. P., Kelly, M., Laumann, T., Miller, K. L., Moeller, S., Petersen, S., Power, J., Salimi-Khorshidi, G., Snyder, A. Z., Vu, A. T., … Glasser, M. F. (2013). Resting-state fMRI in the Human Connectome Project. NeuroImage, 80, 144–168. 10.1016/J.NEUROIMAGE.2013.05.039

60. Snoek, L., Miletić, S., & Scholte, H. S. (2019). How to control for confounds in decoding analyses of neuroimaging data. NeuroImage, 184, 741–760. 10.1016/J.NEUROIMAGE.2018.09.074

61. Stanley, M. L., Moussa, M. N., Paolini, B. M., Lyday, R. G., Burdette, J. H., & Laurienti, P. J. (2013). Defining nodes in complex brain networks. Frontiers in Computational Neuroscience, 7, 169. 10.3389/fncom.2013.00169

62. Uğurbil, K., Xu, J., Auerbach, E. J., Moeller, S., Vu, A. T., Duarte-Carvajalino, J. M., Lenglet, C., Wu, X., Schmitter, S., Van de Moortele, P. F., Strupp, J., Sapiro, G., De Martino, F., Wang, D., Harel, N., Garwood, M., Chen, L., Feinberg, D. A., Smith, S. M., … Yacoub, E. (2013). Pushing spatial and temporal resolution for functional and diffusion MRI in the Human Connectome Project. NeuroImage, 80, 80–104. 10.1016/j.neuroimage.2013.05.012

63. Van Essen, D. C., Smith, S. M., Barch, D. M., Behrens, T. E. J., Yacoub, E., & Ugurbil, K. (2013). The WU-Minn Human Connectome Project: An overview. NeuroImage, 80, 62–79. 10.1016/j.neuroimage.2013.05.041

64. Waller, L., Walter, H., Kruschwitz, J. D., Reuter, L., Müller, S., Erk, S., & Veer, I. M. (2017). Evaluating the replicability, specificity, and generalizability of connectome fingerprints. NeuroImage, 158, 371–377. 10.1016/J.NEUROIMAGE.2017.07.016

65. Yamada, T., Hashimoto, R.-I., Yahata, N., Ichikawa, N., Yoshihara, Y., Okamoto, Y., Kato, N., Takahashi, H., & Kawato, M. (2017). Resting-State Functional Connectivity-Based Biomarkers and Functional MRI-Based Neurofeedback for Psychiatric Disorders: A Challenge for Developing Theranostic Biomarkers. The International Journal of Neuropsychopharmacology, 20(10), 769–781. 10.1093/ijnp/pyx059

66. Yan, C. G., Cheung, B., Kelly, C., Colcombe, S., Craddock, R. C., Martino, A. D., Li, Q., Zuo, X. N., Castellanos, F. X., & Milham, M. P. (2013). A comprehensive assessment of regional variation in the impact of head micromovements on functional connectomics. NeuroImage, 76, 183–201. 10.1016/J.NEUROIMAGE.2013.03.004

